# Sirtuin2 blockade inhibits replication of Human Immunodeficiency Virus-1 and *Mycobacterium tuberculosis* in macrophages and humanized mice

**DOI:** 10.1101/2024.10.27.620499

**Authors:** Vipul. K. Singh, Abhishek Mishra, Khangy Truong, Jose Alejandro Bohorquez, Suman Sharma, Arshad Khan, Franz Bracher, Kangling Zhang, Janice Endsley, Mark Endsley, Andrew P. Rice, Jason T. Kimata, Guohua Yi, Chinnaswamy Jagannath

## Abstract

Coinfections with *Mycobacterium tuberculosis* (Mtb) and HIV-1 present a critical health challenge and require treatment for survival. We found that human M1 macrophages inhibit Mtb growth, while M2 macrophages, characterized by elevated Sirt2 expression, permit Mtb growth. Further, we found that HIV-1 augmented Sirt2 gene expression in MФs. Therefore, we explored the therapeutic potential of sirtuin-modulating drugs in MФs. Sirtinol, a Sirt2 inhibitor, significantly reduced HIV-1 growth in M0, M1, and M2-MФs by >1 log10 over 7 days. Conversely, individual doses of resveratrol and SRT1460, which activate Sirt1, did not affect HIV-1. However, their combination showed a strong synergistic inhibition of HIV-1. The combination of sirtinol with resveratrol was neither synergistic nor antagonistic. Sirtinol upregulated *iNOS* and *ATG5* mRNA in HIV-1 infected MФs in a phenotype-dependent manner. In a humanized mouse model (Hu-NSG-SGM3) co-infected with Mtb H37Rv and the HIV-1 BAL strain, treatment with sirtinol alone, or in combination with combination antiretroviral therapy (cART), showed promising results; Sirtinol alone reduced Mtb growth, while its combination with cART effectively inhibited HIV-1 replication in the organs. We propose that Sirt2 blockade and Sirt1-activation represent a novel dual therapeutic strategy for treating HIV-1 and Mtb coinfections.

## INTRODUCTION

Tuberculosis is the second leading cause of death from infectious diseases, claiming ∼1.2 million lives yearly. An unusual feature of tuberculosis is that the causative pathogen, *Mycobacterium tuberculosis* (Mtb), can either rapidly progress to active lung infection in ∼10% of infected individuals or remain dormant for decades in ∼90%. HIV-1 is a similarly devastating infectious agent. Importantly, one-fourth of the population is latently infected with Mtb (LTBI) and these individuals serve as a reservoir for potential TB reactivation. This risk is exacerbated as HIV-1 progressively depletes CD4 T cells, which are critical for anti-TB immunity. Curiously, HIV-1-infected individuals remain susceptible to TB even when CD4 T cell counts are within normal ranges, suggesting that myeloid/macrophages (MΦ) may play a role during TB/HIV-1 coinfection. Despite the hostile environment of MΦs, Mtb can survive, and HIV-1 can replicate within them, often providing a latent reservoir (1, 2). The lethal nature of untreated coinfections underscores the urgent need for a better understanding of the molecular interplay between Mtb and HIV-1 within MΦs.

Most studies involving Mtb and HIV-1 have used undifferentiated, naïve MΦs. We recently reported functional heterogeneity between human M1 and M2-MΦs compared to naïve MΦs (M0-MΦs) and revealed differences in pro-inflammatory and anti-inflammatory cytokine production correlating with different transcriptomes (3). IFN-γ-stimulated M1-MΦs eliminate Mtb through autophagy and nitric oxide (NO)-dependent mechanisms, while IL-4-stimulated M2-MΦs permit Mtb growth due to decreased autophagy and NO levels. Importantly, Mtb-infected M1 and M2-MΦs exhibit differential induction of NAD^+^ dependent Sirtuin-type histone deacetylases. Blockade of Sirtuin2 (Sirt2) using Sirtinol polarizes naïve or M2-MΦs to M1-MΦs, which are capable of eliminating Mtb. Intriguingly, Sirt2 blockade also inhibits HIV-1 growth in human MΦs, irrespective of their M0, M1, or M2 phenotype. Thus, Sirtuins appear to be critical epigenetic modifiers of the early interactions between Mtb and HIV-1 in MΦs.

Many studies indicate that Mtb and HIV-1 exacerbate each other’s effects; therefore, we hypothesized that Sirt2 activation by Mtb impairs MΦ function, thereby facilitating HIV-1 replication. Intriguingly, the HIV-1 transactivator protein, Tat, binds to and inhibits Sirt1 histone deacetylase, increasing the acetylation of the NF-kB p50/p65 subunit, which leads to T cell hyperactivation and increased HIV-1 replication (4),(5). Therefore, prior infection with HIV-1 may reduce Sirt1 levels in MΦs, increasing Mtb growth (6) (7). Multiple sirtuin inhibitor drugs have been evaluated for chemotherapy (8). In this study, we report that Sirt2 inhibits the replication of HIV1 in human MΦs and is a promising target for developing immuno<u>c</u>hemotherapy (ICT) for treating coinfections.

## RESULTS

### Sirtuin2 blockade inhibits Mtb growth in human macrophages and mice (Fig. 1)

We recently reported that human MΦs exhibit differential responses to Mtb infection, influenced by sirtuin expression levels. Specifically, in IFN-γ (M1) and IL-4 (M2) pre-programmed and rested MΦs that were subsequently infected with Mtb, we found that Sirt5**^hi^** M1-MΦs restricted Mtb through autophagy and NO production. In contrast, Sirt2**^hi^** M2-MΦs permitted Mtb growth due to reduced autophagy and NO levels (3). Mtb infection upregulated transcripts for additional sirtuins in both M1 and M2-MΦs (**Fig. 1A**). Further, infecting naïve M0-MΦs using virulent Mtb (but not the attenuated BCG vaccine) progressively increased Sirt2 gene expression (**Fig. 1B**). PCR studies revealed elevated Sirt2 levels in PBMCs from patients with active TB compared to healthy contacts, who showed an upregulation of Sirt5 (3). These findings suggest that Sirt2 is a druggable target for Mtb control in human MΦs. **Fig. 1C** shows that Sirt2 blockade using sirtinol, but not the Sirt5 inhibitor balsalazide, inhibited Mtb growth in naïve MΦs. Sirt1 inhibition was also ineffective, consistent with reports that Sirt1activation might suppress Mtb growth in MΦs (9). To determine the role of Sirt2 *in vivo*, C57BL/6 mice were aerosol-infected using Mtb Erdman, followed by sirtinol treatment. Sirtinol treatment significantly reduced bacterial growth in the lungs (**Fig. 1D**). Together, these data indicate that targeting Sirt2 with sirtinol is an effective strategy for controlling tuberculosis in both MΦs and mouse models.

**FIGURE 1:**
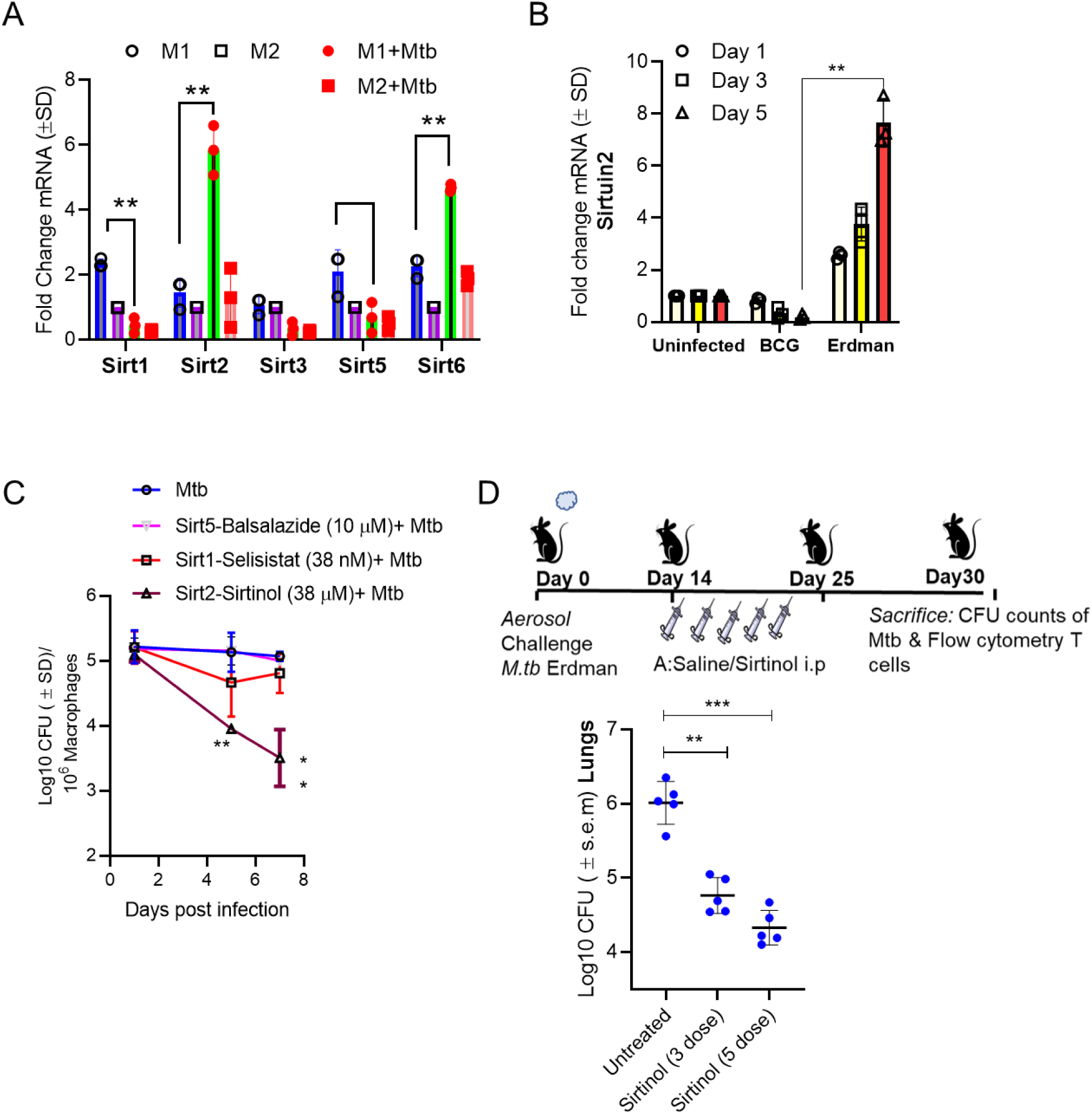
Sirtuin2 histone deacetylase regulates *Mycobacterium tuberculosis* growth in human macrophages and C57BL/6 mice. CD14+ bead-purified macrophages derived from healthy donors (MФs; pooled from 3 individuals) were either left naïve (M0) or differentiated into M1-MФs with IFN-γ or M2-MФs with IL-4, then rested and processed as follows. **A:** M1 and M2-MФs were infected with Mtb H37Rv (MOI=1) for 4 h, washed, incubated, and Trizol lysates from triplicate wells collected 24 h post-infection were assessed for mRNA expression of sirtuins as indicated (** p< 0.009; two-tailed t-test). **B:** Pooled MФs (n=3 donors per pool; 2 pools) were infected with either the BCG vaccine or Mtb (MOI=1), washed, incubated, and Trizol lysates were tested for mRNA expression of Sirt2 and Sirt5 (** p< 0.009; two-tailed t-test). **C:** Pooled MФs (n=3 donors per pool; 2 pools) infected with Mtb were cultured and then treated with the indicated drugs (sirtinol for Sirt2 inhibition at 38 µM); selisistat for Sirt1 inhibition at 38 nM; balsalazide for Sirt5 inhibition at 25 µM), followed by a colony forming unit (CFU) assay (** to measure bacterial growth (** p< 0.005; one-way ANOVA). **D:** Age and sex-matched C57BL/6 mice (n=5 per group, 4-8 weeks, male + female) were infected with Mtb H37Rv (100 CFU per mouse) using a Glas-Col aerosol chamber. On days indicated days post-infection, mice were injected intraperitoneally with 10 mg/kg body weight Sirtinol. Mice were sacrificed on day 30 post-infection to assess Mtb CFU counts in the lungs (** p< 0.007*** p< 0.006; two-way ANOVA with Tukey’s posttest).

### HIV-1 infection induces sirtuins in human macrophages. (Fig. 2)

Because the HIV-1 transactivator Tat protein binds and inhibits Sirt1, increasing HIV-1 replication (4),(5), we hypothesized that it may also induce Sirt2 in MΦs. Using donor-derived MΦs, we identified that, as early as 18 h post-infection, HIV-1 increased the levels of mRNA transcripts for Sirt1, Sirt2, and Sirt3 in both naïve (M0) and M1-MΦs, whereas M2-MΦs displayed a relatively reduced response to infection (**Fig. 2A**). Sirt1-7 regulate multiple functions in mammalian cells and our previous research has shown that blockade of one sirtuin can enhance the gene expression of others. Given the prominent roles of Sirt1 and Sirt2, we treated HIV-1 infected MΦs with the Sirt1 activator resveratrol and the Sirt2 inhibitor sirtinol. Subsequent PCR analysis of sirtuins revealed that inhibiting Sirt2 with sirtinol resulted in increased expression of Sirt5,6 and 7 in naïve M0-MΦs, suggesting there may be a cross-regulation among sirtuins upon Sirt2 blockade. Sirtuins can also influence autophagy and nitric oxide (NO)-dependent antimicrobial mechanisms in MΦs (10) (11). **Fig. 2B-C** illustrates that Sirt2 blockade upregulated mRNA transcripts for autophagy (*ATG5*) and NO production (*iNOS*) biomarkers in MΦs. However, iNOS protein levels decreased after Sirt2 blockade in HIV infected MΦs for which we have no clear explanation; perhaps HIV-1 interferes with translation of *iNOS* gene (12) (13).

**FIGURE 2:**
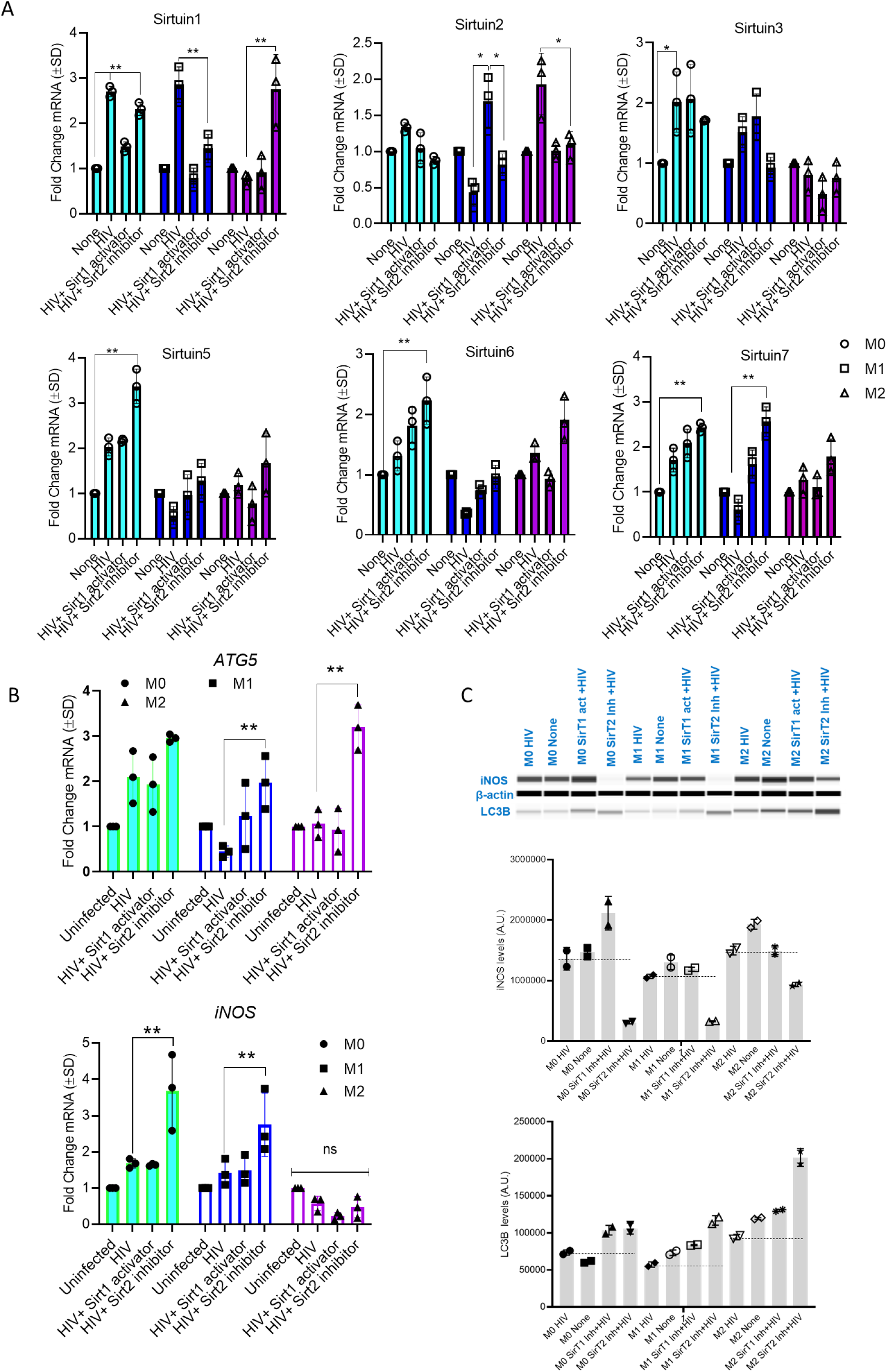
Differential expression of Sirtuin mRNA affects nitric oxide and autophagy in HIV-1 infected human macrophages. **A:** MФs pooled from 3 donors each, prepared as naïve (M0), M1, or M2 as in Fig.1, were infected with HIV-1 (NL-AD8 strain; stock virus concentration of 1 × 10^6 TCID_50_/mL. For infection, 100 µL of the virus was added to achieve a multiplicity of infection of 0.1 (TCID_50_). Trizol lysates were collected on day 3 post-infection. Replicate sets of these samples were treated with a Sirtuin1 activator (Resveratrol, 20 µM) or a Sirtuin2 inhibitor (Sirtinol, 30 µm) and subsequently analyzed via QPCR for the indicated Sirtuins, (*p< 0.006 ** p <0.007; two-tailed t-test). **B:** Trizol lysates from panel A were assayed using primers for human inducible nitric oxide synthase (iNOS) and Autophagy regulating gene 5 (*ATG5*). **C:** Additional replicate sets of MФs from the same batch wells were pooled and analyzed for protein expression using mAbs targeting iNOS and light chain associated with microtubule B (LC3B), a biomarker for autophagy. Western blots were conducted using capillary gel electrophoresis. Densitometry quantitation (average units; AU) is shown (* p< 0.009; t-test).

### Sirtuin2 blockade inhibits HIV-1 replication in human macrophages. (Fig. 3)

Our data indicate that Sirtuins, including Sirt1 and Sirt2, differentially regulate antimicrobial functions and are therapeutic targets for HIV-1 infection. Because treating infection and coinfection is difficult, and often requires multiple drugs, we first examined the effects of Sirt1 activation (due to the interaction of HIV-1 Tat with Sirt1) and Sirt2 blockade on HIV-1 replication. We observed a dose-dependent effect of noncytotoxic doses of Sirt1 activators (resveratrol, 20 μM; SRT1460,3 μM) and inhibitors of Sirt2 (sirtinol, 30 μM), Sirt3 (3-TYP; 377 μM) and Sirt5 (balsalazide; 10 μM) to inhibit HIV-1. When administered as a single drug dose, 24 h post HIV-1 infection, only sirtinol, a Sirt2 inhibitor, showed a significant 2log10 reduction in HIV-1 replication in MΦs (**Fig. 3A**). Conversely, resveratrol (a Sirt1 activator) and balsalazide (a Sirt5 inhibitor) also reduced HIV-1 replication, but to a lesser extent (1log10 between days 3 and 7). In two separate experiments, we observed that HIV-1 replication resumed after it was initially drastically reduced by sirtinol (**Fig. 3A**). Therefore, in three separately conducted experiments using different sets of donor cells, drugs were administered on day 1 post-infection and re-applied on days 3 and 5. Remarkably, sirtinol consistently inhibited HIV1 replication (a 2log10 reduction at day 3 post-infection and 3log10 decrease by days 5 and 7) (**Fig. 3B**). Intriguingly, neither resveratrol nor SRT1460 (Sirt1-activators) alone inhibited HIV-1, although their combined use resulted in a synergistic effect, reducing HIV1 replication by 1-2 log10. A combination of sirtinol and resveratrol was also effective, suggesting that resveratrol is not antagonistic to sirtinol. MΦ viability remained >90% in these assays. Although higher doses of sirtuin-modulating drugs were not evaluated, these data suggest that Sirt1 activation and Sirt2 blockade are effective strategies for controlling HIV-1 replication in MΦs.

**FIGURE 3:**
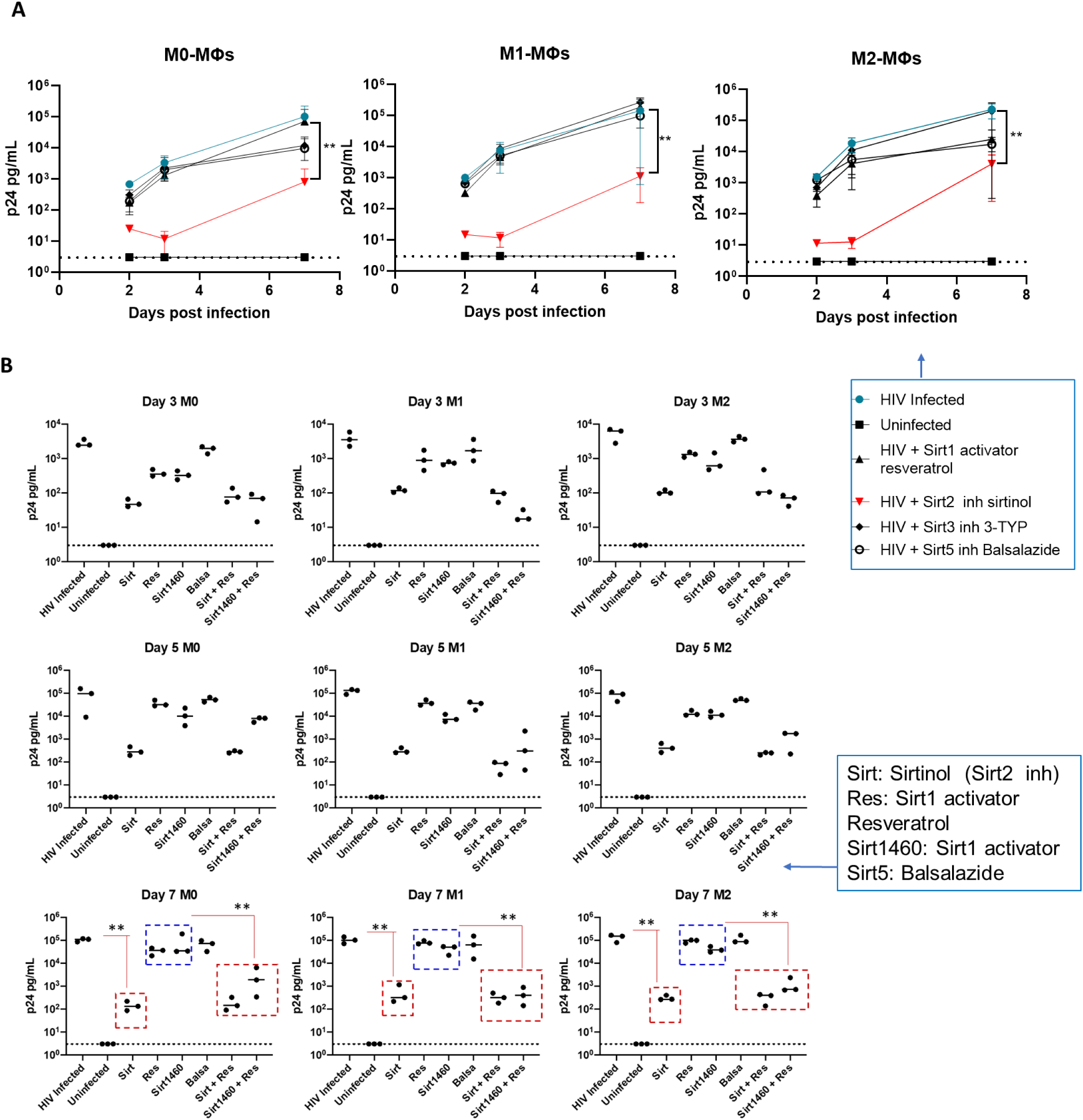
**S**irt1 activation and Sirt2 blockade restrict HIV-1 replication in macrophages. A: Macrophages were infected at a MOI of 0.1 with HIV-1 NLAD8 as described in the materials and methods. After 24 hours, infected cells were treated with the indicated Sirt activators or inhibitors. HIV-1 production was quantified by HIV-1 p24 ELISA at 2, 3, and 7 days post-infection. (** p <0.007; two-tailed t-test). B: Macrophage infections with HIV-1 NLAD8 were performed as in A but the drugs were applied at 24 hours post-infection, and re-applied on days 3 and 5 post-infection *(\*\** p< 0.01; two tailed t-test).

### Sirt2 inhibition with sirtinol and Sirt1 activation with resveratrol increases Rel-A/NF-kB acetylation. (Fig. 4)

A major regulatory mechanism during microbial infection is the acetylation and methylation of histones and enzymes that regulate the expression of immunity-related genes and the polarization of MΦs (14). In the context of HIV-1, NAD-independent HDACs regulate viral gene expression and HDAC inhibitors have been used for therapy. Compounds like entinostat reactivate latent HIV-1 and, when used in combination with the protein kinase c agonist, bryostatin-1, induce maximal HIV-1 protein expression (15). Given the effectiveness of Sirt1 activation and Sirt2 blockade against HIV-1 (**Fig. 3**), we used donor-derived MФs, either as naïve M0 or differentiated into M1 or M2, rested them, and then infected them with HIV-1, following the earlier described drug treatment protocol (**Fig. 3**). Cell lysates collected on day 4 post-drug treatment were analyzed for Rel-A acetylation using multi-OMICS approaches (16). **Fig. 4** shows that both sirtinol and resveratrol increased Rel-A acetylation compared to levels observed in HIV-1-infected MФs alone. Because NF-kB Rel plays both agonistic and antagonistic roles during HIV-1 replication (17), additional investigations are warranted to define the specific anti-HIV-1 effects of Sirt2 inhibition, focusing on the NF-kB complex.

**FIGURE 4:**
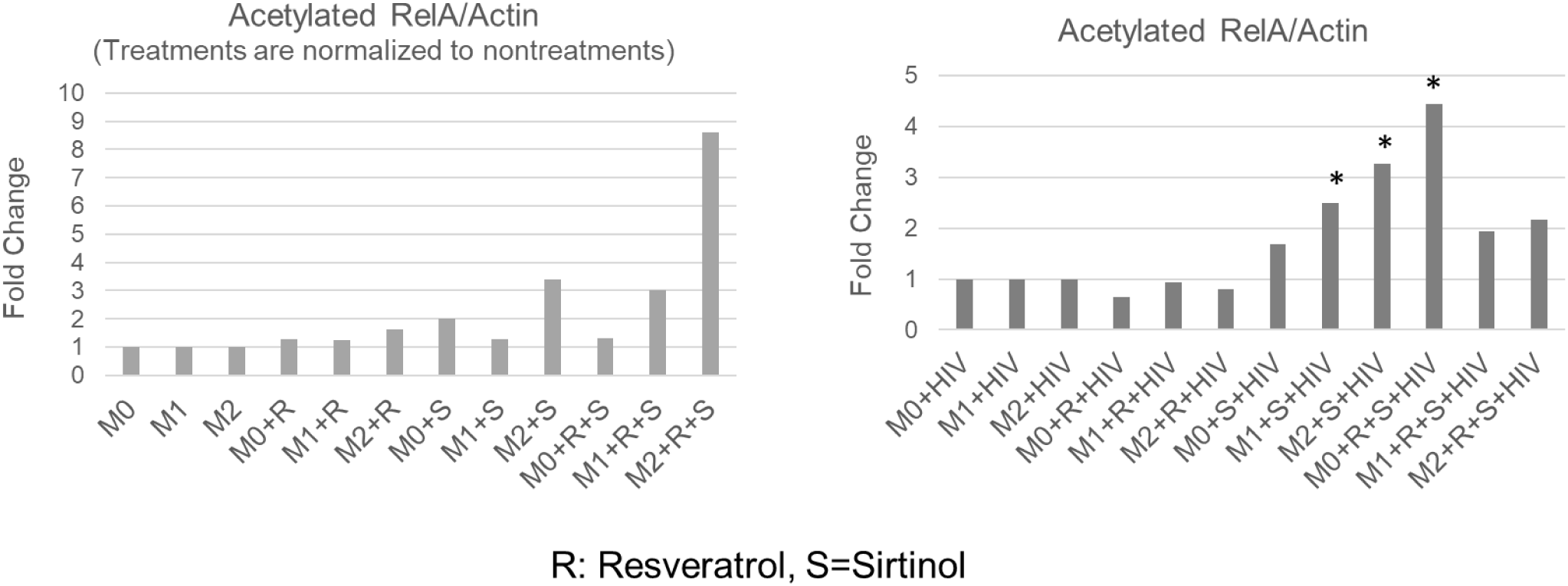
Sirt2 blockade with sirtinol and Sirt1 activation with resveratrol increase Rel-A/NF-kB acetylation. MФs derived from human donors were either left naïve (M0) or differentiated into M1 or M2 phenotypes, followed by HIV-1 infection as described in Fig. 3A. Following infection, cells were treated with sirtuin modulators, including Sirtinol (30 µM) alone or Sirtinol and resveratrol (30 µM). Cell lysates were collected on day 4 post-drug addition and analyzed for Rel-A acetylation using OMICS Left: Uninfected MФs. Right: HIV-infected M0, M1, and M2-MФs. (* p< 0.01; vs. HIV alone groups, t-test). *Supplemental* Fig.4 illustrates data analysis.

### Sirt2 blockade using sirtinol suppresses both HIV and Mtb replication during coinfection of MФs (Fig. 5)

Both HIV-1 and Mtb can coinfect MΦs, and our findings indicate that Sirt2 blockade suppresses replication of both pathogens (**Figs. 1 and 3**). Thus, we investigated whether sirtinol could simultaneously inhibit HIV/Mtb replication in MΦs. **Fig. 5** shows that M1 and M2-MФs (rested and subsequently infected with HIV) from four healthy donors exhibited similar levels of HIV-1 RNA loads three days post-infection, regardless of M1 or M2 phenotype or treatment regimen (sirtinol, cART, or sirtinol/cART). Seven days post-infection, a significant reduction in viral RNA levels was observed in HIV-infected M2-MФs treated with sirtinol compared to untreated controls. The viral RNA load of the sirtinol-treated M2-MФs was similar to levels observed following either single cART or combined cART/sirtinol treatment. Co-infection with Mtb did not significantly affect HIV-1 replication, as both M1 and M2-MФs showed similar viral loads in both the HIV-1 only and HIV/Mtb confection groups (**Fig. 5A**). Of note, Mtb bacterial load was also significantly reduced seven days post-infection in HIV/Mtb co-infected M2-MФs after sirtinol treatment (**Fig. 5B**). Reduced Mtb load was observed only in sirtinol-treated M2-MФs, with or without additional cART treatment. Conversely, M2-MФs treated only with cART showed Mtb bacterial loads comparable to those of untreated MФs, whether infected with Mtb alone or with HIV/Mtb (**Fig. 5B**).

**FIGURE 5:**
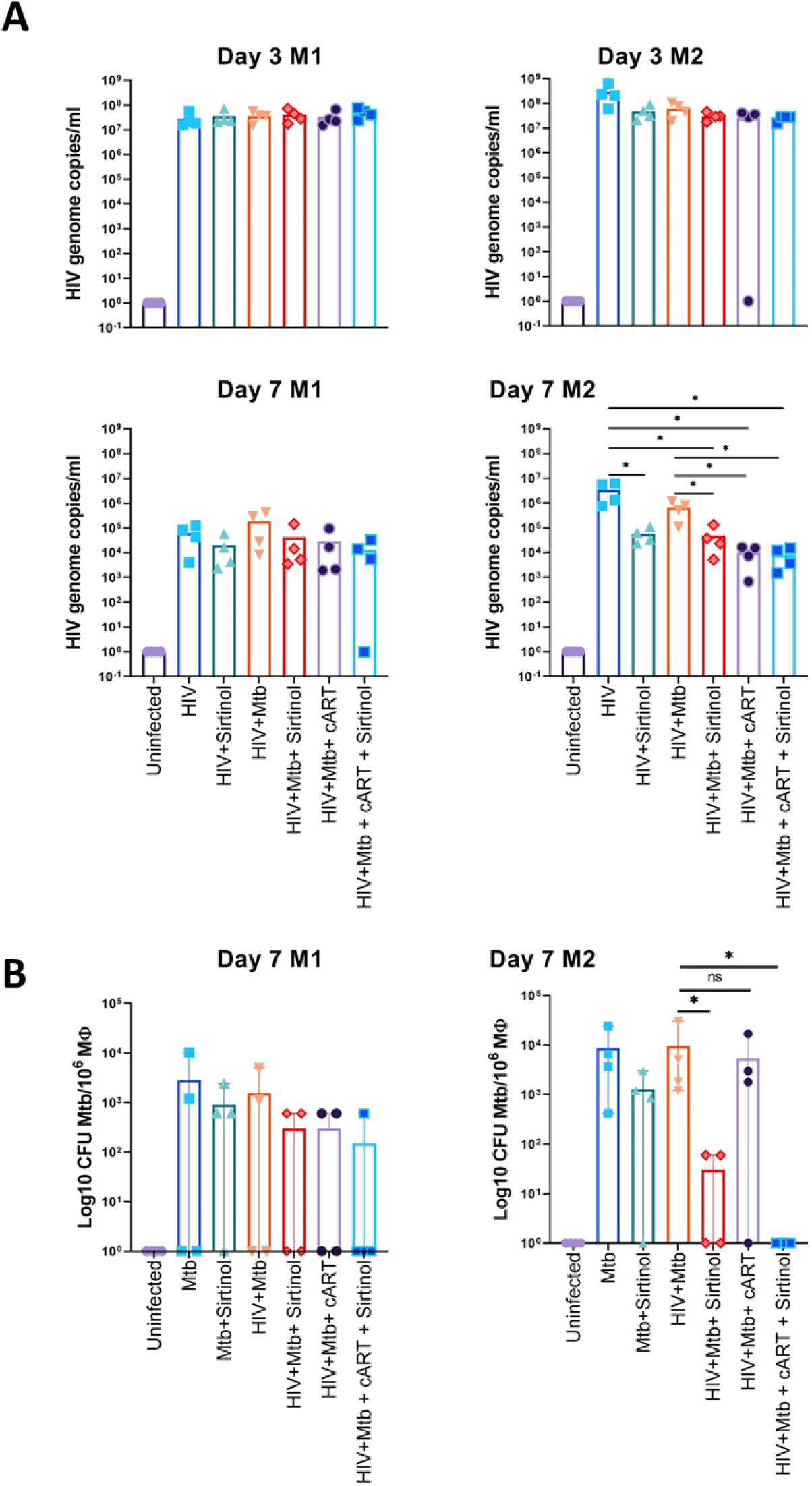
Sirt2 blockade using sirtinol inhibits both HIV and Mtb replication in co-infected macrophages. MФs from 4 donors were differentiated into M1 and M2 phenotypes before being infected with either HIV, Mtb, or both pathogens together. Then, various treatments were applied as indicated. **A:** Viral loads for HIV were assessed from supernatants collected on days 3 and 7; **B:** Mtb CFUs were quantified from MФ lysates on day 7 (* p< 0.01; paired t-test).

### The Hu-NSG-SGM3 mouse model permits replication of both HIV-1 and Mtb infections. (Fig. 6)

To establish an animal model for HIV/Mtb coinfection, we first infected humanized NSG-SGM3 mice with Mtb using the aerosol route. In coinfected groups, this was followed by intraperitoneal infection with HIV-1. Mice receiving either Mtb alone or HIV-1 alone, and mice left uninfected were used as controls. Post-infection, HIV-1-infected mice exhibited a decreased CD4/CD8 T-cell ratio compared to uninfected controls (**Fig. 6A**). Of note, HIV-1 RNA was detected in the humanized mice starting at 28 days post-infection (dpi), with a viral RNA load between 10^7^ and 10^8^ genome copies/mL of serum, with an increase observed by 35 dpi (**Fig. 6B**). Mtb infection was confirmed with detection of a bacillary load of around 100 colony forming units (CFU) in the lungs of control animals euthanized one day post-infection (**Fig. 6C**). At the end of the study (35 dpi), lung Mtb loads ranged from 10^5^ to 10^6^ CFU in the Mtb single-infection animals and from 10^4^ to 10^6^ CFU in the HIV/Mtb co-infected mice, though no statistical differences were found between these groups. Pulmonary function tests in these mice demonstrated significantly reduced lung volume and compliance, as well as increased elastance and resistance in the HIV/Mtb coinfected mice compared to uninfected controls (**Fig. 6D**). CT scans also showed multiple high-density areas in the lungs of Mtb-infected animals, with and without HIV-1 coinfection (**Fig. 6E**), indicating inflammation and pathological changes. Although Mtb was also found in the spleens, the average CFU counts were lower than in the lungs; loads of 1.5×10^4^ and 1.8×10^5^ in the Mtb alone and HIV/Mtb coinfection groups, respectively, suggested that HIV-1 coinfection elevated spleen Mtb loads.

**FIGURE 6:**
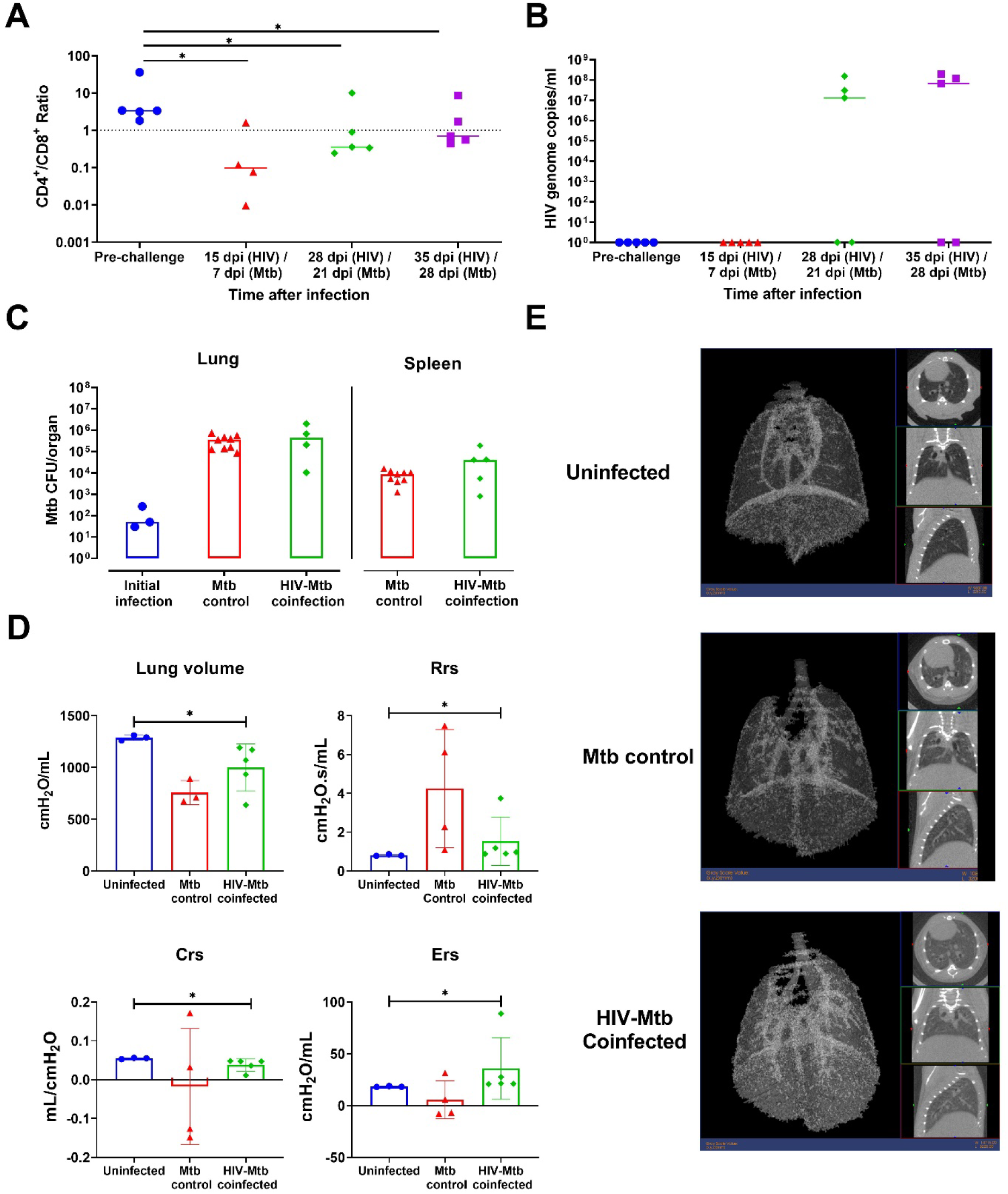
Humanized mice (Hu-NSG-SGM3) allow HIV-1 and Mtb replication during coinfection. Hu-NSG-SGM3 mice were initially infected intraperitoneally with 10^5 TCID50 of the HIV-1 BAL strain, (for groups designated as HIV-alone and HIV/Mtb coinfection). Seven days post-HIV infection, an aerosolized dose of Mtb H37Rv (100 CFU per mouse lung) was administered using a Madison chamber, as previously described by us (26) in both Mtb-only and HIV/Mtb coinfection groups. Day 0 implantation of Mtb was verified through lung cultures from sacrificed mice. **A and B:** CD4+ T cell levels (A) and HIV viral load (B) were monitored across different time points. **C:** Mtb bacillary load in lung and spleen tissues was assessed at the study’s conclusion. **D:** Pulmonary function was evaluated by measuring lung volume and respiratory resistance (Rs), elasticity (Ers), and compliance (Crs). **E:** CT scans were performed, and one representative figure is shown for each group. The left panels show the 3D images, with white areas indicating high-density scans (e.g., tissues) and black areas indicating low-density scans (e.g., air). The three smaller figures for each mouse show various scan angles per mouse. Unpaired multiple t-tests were used for statistical analysis. ns: p ≥ 0.05, no significant difference; *: p< 0.05; ** p< 0.01; and *** p< 0.001.

### Effect of Sirt2 blockade using sirtinol treatment in humanized mice coinfected with HIV/Mtb. (Fig. 7)

To determine the effect of sirtinol on HIV/Mtb coinfection, humanized mice were treated with sirtinol intraperitoneally (**Fig. 7A**). No significant differences in HIV RNA levels were detected in the HIV/Mtb co-infected mice after sirtinol treatment when compared to untreated (but HIV/Mtb coinfected) mice, though one animal exhibited undetectable levels of viral RNA, which had previously been positive by HIV RT-qPCR. However, mice receiving cART alone or cART+Sirtinol displayed a significant reduction in viral RNA loads (**Fig. 7B**).

**FIGURE 7:**
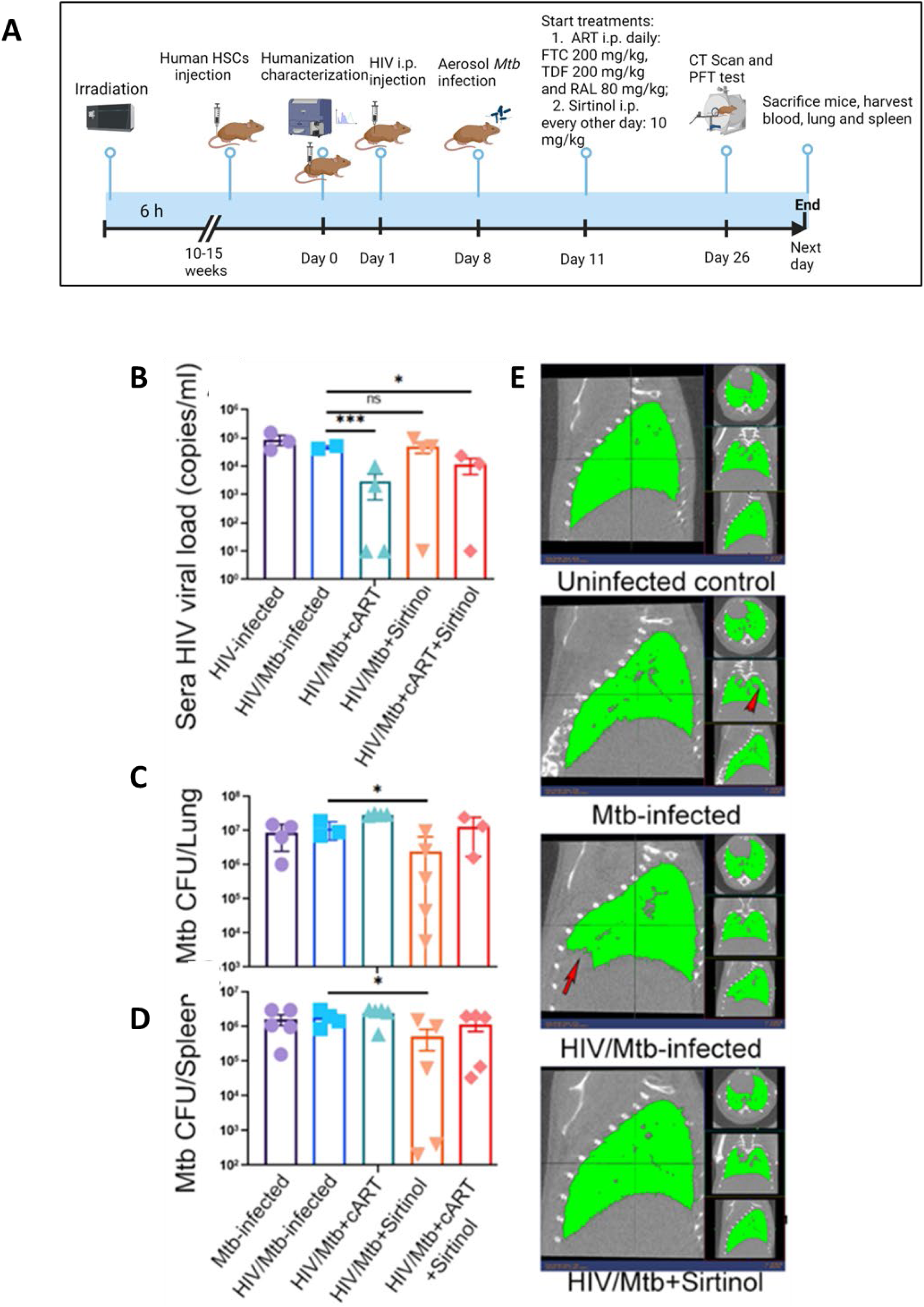
Sirtinol inhibits replication of both HIV/Mtb in Hu-NSG-SGM3 mice. **A:** Protocol for coinfection and treatment. Humanized NSG-SGM3 mice were infected with HIV and/or Mtb, followed by administration of various treatments as indicated. cART was administered daily while sirtinol (10 mg/kg body weight) was given every other day (totaling 12 doses over 24 days). Post-treatment, mice were sacrificed for CT scans, followed by collection of sera and tissue samples to determine HIV viral loads and Mtb CFU counts. **B:** Serum HIV viral load measurements. **C and D:** Mtb CFU counts within the lung (C) and spleen (D). **E:** the lung CT scans, with gray areas within the functionally illustrated lungs (green) depicting inflammation or pathological changes (such as lesions, etc.). The obvious lesions visible to the naked eye are marked with red arrows. Unpaired multiple t-tests were used for statistical analysis. ns: p≥ 0.05, no significant difference; * P< 0.05; ** p< 0.01; and *** p< 0.001.

The Mtb bacterial load in the sirtinol-treated mice was significantly lower than that in untreated mice, for both lung and spleen samples (**Figs. 7B-7C**). No significant differences in Mtb load were found in mice treated with cART alone or those receiving cART+sirtinol therapy. CT scans showed extensive lung lesions in Mtb-infected animals, with or without HIV co-infection. Conversely, lungs from sirtinol-treated mice showed smaller lesions that were comparable to those of uninfected mice (**Fig. 7D**).

## DISCUSSION AND CONCLUSIONS

*M. tuberculosis* and HIV-1 cause serious infections that are difficult to control and require complex, multi-drug treatment regimens over extended durations. Both pathogens persist for decades in the human body and can be reactivated from a latent state. While HIV-1 primarily targets CD4 T cells, it can also infect MФs (18), which recent studies have identified as a source for the virus in individuals who are unresponsive to cART, thereby rejuvenating research into this relatively unexplored niche (18). Similarly, Mtb can infect many cell types but predominantly replicates and persists within MФs. For example, within multi-nucleate giant cells of TB granulomas, which arise through the fusion of MФs. Indeed, host-directed therapies for TB mostly center around the activation of MФs. Given the emerging importance of MФs during HIV/Mtb coinfection, we present a novel paradigm that Sirt2 and Sirt1 modulation can influence the outcome of coinfection.

In our recent study, we found that human MФs exhibit functional heterogeneity. Specifically, IFN-γ preconditioned M1-MФs effectively controlled Mtb growth through autophagy and NO production. In contrast, IL-4 preconditioned M2-MФs, lacking these antimicrobial mechanisms, permitted increased Mtb growth (3). These phenotypic differences were reflected in distinct transcriptomic profiles, with M1-MФs showing stronger Sirt5 expression and M2-MФs showing marked Sirt2 upregulation (19). Notably, Sirt2 blockade with sirtinol not only enhanced Mtb clearance in MФs and C57BL/6 mice (**Fig.1**), but also facilitated a phenotypic switch from M2 to M1, as evidenced by dramatic changes in proteomics and anti-microbial function (Mishra *et al.* under submission). Unexpectedly, HIV-1 infection also upregulated Sirt2 in addition to Sirt1 in MФs (**Fig. 2**). We have recently described that Sirtuins play a major role in regulating histone modifications that affect MФ polarization and antimicrobial function (16) (20) (Zhang, Mishra, Jagannath, *in press*). Herein, we demonstrate a novel immunoregulatory role for Sirtuins during both standalone and coinfections with Mtb and HIV-1 in MФs.

Though the HIV-1 derived Tat protein enhances viral replication by binding to Sirt1 in T cells (4),(5), we found that Sirt2 blockade dramatically inhibits HIV1 replication in MФs (**Fig. 3**). Although Sirt1 activation with resveratrol alone did not inhibit HIV-1, its combination with SRT1460 significantly reduced HIV-1 replication in MФs. Because we used only two doses of these drugs on day 2 and 5 to inhibit HIV-I *ex vivo,* and because multiple dosing during chemotherapy is feasible, we propose that a longer duration of treatment with multiple or higher doses could offer a more effective treatment regimen.

Similar to Mtb, HIV-1 also induces mRNA transcripts for Sirt1, Sirt2, Sirt3 and detectable levels of Sirt5, Sirt 6, and Sirt7. It is possible that these latter sirtuins also play a role during the progression of HIV-1 pathogenesis. Sirt2 blockade increased transcripts for Sirt5, Sirt 6, and Sirt7 in HIV-1 infected MФs (**Fig. 3**), suggesting there is a cross-regulatory mechanism among sirtuins that warrants additional investigation. Sirtuins are histone deacetylases that epigenetically modify gene expression. Interestingly, it has been shown that HIV-1 Tat binding to Sirt1 inhibits its activity thereby increasing acetylation of the NF-kB p50/p65 subunit, which hyperactivates T cells and supports HIV-1 replication (4),(5). This may also explain the increase in acetylation of Rel A in HIV-1 infected macrophages (Fig. 4). Sirt1 is also critical for Tat-mediated transactivation of HIV-1. The paradoxical observation underscores the agonistic and antagonistic functions of NF-kB:Rel and Sirt1 in the context of HIV-1 replication along and emphasizes the need for additional research(17).

Here we show that sirtinol reduces the growth of HIV-1 in coinfected MФs and in the organs of humanized mice. The inhibitory effect of sirtinol on HIV-1 was most evident in in vitro MФs infections. However, sirtinol alone did not appear to inhibit HIV-1 effectively, as it achieved significant suppression only when combined with cART. Much of the virus quantified in serum likely originated from CD4+ T cells, potentially masking the inhibitory effect of sirtinol on HIV-1 produced by macrophages in organs, such as the lungs. Further analysis of HIV-1 cell-associated RNA in lung tissue is warranted to determine the impact of sirtinol on HIV-1 expression and its potential association with lesion reduction. Additionally, longer treatment durations with cART and sirtinol may provide additional insights into whether maximal suppression of HIV-1 can further reduce Mtb burden and lesion severity.

## ACKNOWLEDGEMENTS

This work was supported in part by the Texas Developmental Center for AIDS Research, an NIH funded program (P30 AI161943); NIH AI116167 to JTK, NIH AI161015 to CJ, and NIH AI150550 to GY. FB was supported by Deutsche Forschungsgemeinschaft (DFG) with funds from SFB1309 (Chemical Biology of Epigenetic Modifications), project-ID: 325871075– SFB1309.

## AUTHOR CONTRIBUTIONS

Authors performed experiments as follows: AM (Fig.1,2), VKS,KT, SS, AK (Fig.3), KZ (Fig.4), FB provided reagents, JAB, GY performed Humice (Figs. 5,6,7), JJ, ME, APR, JK, GY and CJ overall designing, data analysis and writing manuscript. Authors are grateful for Dr. Heather McConnell for editing the manuscript.

## MATERIALS AND METHODS

### Macrophages

Healthy donor derived MФs were purified from PBMC using CD14 magnetic beads their differentiation into M0 (naïve), M1 and M2-MФs has been described by us (3). MФs were forst infected using Mtb (MOI=1) or HIV-1 followed by treatment with Sirtuin (1/2/3/5) inhibitors or activators and growth assayed over 7 days using Mtb CFU counts or p24 ELISA. Sirtuin modulators were also combined with anti-retroviral drugs (ART) where indicated. MФ lysates were assayed for Sirtuins, *iNOS* and *ATG5/7* gene expression and western blots performed as described by us (3).

### Cell lines and infectious HIV-1

The R5 macrophage-tropic plasmid proviral clone, pNL(AD8) (NLAD8), and TZM-bl cell line were obtained from the NIH AIDS Reagent Program (21–23). 293T cells were obtained from the American Type Culture Collection (ATCC, CRL-3216). Both the 293T and TZM-bl cell lines were maintained in complete Dulbecco’s modified Eagle’s medium (DMEM) (i.e., high-glucose (4.5 g/L) DMEM (Corning, 10-0130-CV) supplemented with 10% fetal bovine serum heat-inactivated at 56°C for 30 minutes, 2 mM L-glutamine, 1 mM sodium pyruvate, 100 U/ml penicillin, 100 μg/ml streptomycin). Stocks of infectious HIV-1 NLAD8 were produced by transfection of 293T with pNL(AD8), and the infectious titers were determined by limiting dilution using TZM-bl cells to score infection by luciferase assay, as we have previous described (24).

### Macrophage infection and HIV-1 p24 ELISA

1×10^6 naïve, IFN-gamma, or IL-4 pre-programmed macrophages in 24 well plates were infected with HIV-1 NLAD8 strain at a MOI of 0.1 for 24 hours. After 24 hours, cells were washed once using 1 mL of Dulbecco’s phosphate - buffered saline (PBS) without calcium and magnesium (Corning, 21-031-CV) to remove unattached virions. 1 mL of fresh media was replaced and supernatants were collected to quantify HIV-1 p24 antigen by antigen capture ELISA (Advanced Bioscience Laboratories, #5447), according to the manufacturers protocol. Absorbance was determined using a universal micro-plate reader (BIO-TEK instruments ELx800). Test sample HIV-1 p24 concentrations were calculated from a standard curve by linear regression analysis. Significance were determined by the appropriate statistical analysis in GraphPad Prism.

### Coinfection procedures

#### Bacterial and viral strains

Mtb H37Rv was obtained from BEI Resources and propagated in the biosafety level 3 (BSL-3) facilities at the University of Texas Health Science Center at Tyler (UTHSCT). Mtb was cultured in 7H9 media with 10% ADC following standard Mtb culture procedures30. After 7 days of growth, the bacteria were collected and subjected to sonication three times, at an amplitude of 38%, for 10 seconds/each, with a 5-second interval, followed by low-speed centrifugation (1,100 RPM). Bacteria were diluted to an optical density (OD) value of ≈ 1 in sterile NaCl 0.9% and aliquots were made and frozen at −80 °C to be used as inoculum. Two weeks later, one aliquot was thawed, and the bacterial load was evaluated by plating ten-fold serial dilutions in 7H10 agar, supplemented with OADC. After 2-3 weeks of incubation, the colony forming units (CFU)/ aliquot were calculated.

#### HIV-1 BAL strain

This was obtained from NIH AIDS Reagent Program, also prepared in the BSL-3 facilities at UTHSCT, following standard procedures 31. Briefly, frozen human PBMCs (STEMCELL Technologies, Vancouver, Canada) were thawed and seeded in a 75 cm2 flask at a concentration of 5 × 106 cells/ml in RPMI 1640 media (Corning Inc., Corning, NY) supplemented with 10% fetal bovine serum (FBS), 1% penicillin/streptomycin, 1 µg/ml of PHA and 2 µg/ml polybrene (MilliporeSigma, Burlington, MA). After 3 days of stimulation, 4 × 107 cells were centrifuged and infected with 1.5 ml of HIV-1 in two adsorption cycles. Following the second adsorption cycle, the cells were seeded in two 75 cm2 flasks with 30 ml of media supplemented with FBS, antibiotics, and human IL-2 (20 Units/ml). Cell culture supernatant was collected every three days, with fresh media being added, until day 21 of culture and stored at −80 °C. Animal experiment design

#### Humanized mice

(25) All animal procedures were approved by the UTHSCT Institutional Animal Care and Use Committee (IACUC). NOD.Cg-Prkdcscid Il2rgtm1Wjl Tg(CMV-IL3,CSF2,KITLG)1Eav/MloySzJ (NSG-SGM3) mice were purchased from The Jackson laboratory (Bar Harbor, ME) and bred in the Vivarium facilities at UTHSCT. Pups were weaned at 21 days after birth and, 1-3 weeks after that, they were irradiated at a dose of 100 cgy/mouse, followed by intravenous injection with 2 × 105 CD34+ stem cells/mouse at 12 h post-irradiation. Humanization was monitored starting at 12 weeks after stem cell transplantation and again at 14 and 16 weeks. For this purpose, blood was drawn from the submandibular vein (100-150 µl, based on animal weight) and PBMCs were collected through density gradient centrifugation using Ficoll Paque (Cytiva, Marlborough, MA). After erythrocyte lysis, the PBMC from each animal were stained for human (hu) and mouse (mo) hematopoietic cell surface marker (CD45+), as well as lymphocytic and myeloid markers. Animals that showed a positive huCD45+/moCD45+ ratio, accompanied by differentiation of various immune cell populations, were selected for experimental infection.

Mice were randomly divided into four experimental groups: Uninfected (n=5), HIV-infected (n=8), Mtb-infected (n=8) and HIV/Mtb co-infected (n=7). Mtb infection was performed using aerosolized Mtb H37Rv through a Madison chamber, as previously described32, using an infection dose of 100 CFU/lung. Three additional mice were included in the Madison chamber at the time of infection and were euthanized 24 hours after infection. The lung sample was collected, macerated and cultured in 7H10 agar, to confirm the initial bacterial load of infection33.

One day after Mtb infection, the mice from the single HIV infection and HIV/Mtb co-infection groups were subjected to intraperitoneal (IP) inoculation with 105 TCID50 of HIV-1 BAL strain. Blood samples from all experimental groups were collected on the day of infection and at 15-, 28- and 35-days post infection (dpi). Serum samples from all the animals were separated and stored at −80 °C until further use. PBMCs were isolated and stained for flow cytometry analysis. At 35 dpi, the animals were terminally anesthetized, using a Ketamine/Xylazine mixture, in order to perform computed tomography (CT) scan and pulmonary function (PF) tests. Afterwards, the animals were euthanized and whole blood samples were collected through cardiac punction. During necropsy, lung and spleen samples were collected and macerated through a 70 μM cell strainer (Thermo scientific) in a final volume of 2 ml of PBS. Serial ten-fold dilutions of the organ macerates were plated in 7H10 agar, supplemented with OADC, to assess the bacterial load. The remaining volume of lung and spleen macerates were stored at −80 °C for further analysis.

For each experimental group, lung sample from one animal was selected for histopathological analysis and, therefore, not subjected to maceration and bacterial culture. Lungs were filled with 10% formalin, before being removed from the animal, and stored in the same media after the necropsy34, 35. Sample processing and Hematoxylin-Eosin (HE) staining was carried out at the histopathology core of UT southwestern.

#### CT scan and PF testing

Mice were IP-injected with ketamine/xylazine (100 mg/kg Ketamine, 20 mg/kg Xylazine). Once the correct anesthetic plane was achieved, the mice were intubated with a sterile, 20-gauge intravenous cannula through the vocal cords into the trachea. Following intubation, anesthesia was maintained using isoflurane._Elastance (Ers), compliance (Crs), and total lung resistance (Rrs) was assessed for each mouse through the snapshot perturbation method, as previously described36. Measurements were performed in triplicates for each animal, using the FlexiVent system (SCIREQ, Tempe, AZ), with a tidal volume of 30 mL/kg at a frequency of 150 breaths/min against 2–3 cm H2O positive end-expiratory pressure. After PF testing, the mice were subjected to CT scans for the measurements of lung volume, using the Explore Locus Micro-CT Scanner (General Electric, GE Healthcare, Wauwatosa, WI). CT scans were performed during full inspiration and at a resolution of 93 μm. Lung volumes were calculated from lung renditions collected at full inspiration. Microview software 2.2 (http://microview.sourceforge.net) was used to analyze lung volumes and render three-dimensional images.

#### RNA extraction and RT-qPCR

Serum samples from all experimental groups were extracted using the NucleoSpin RNA isolation kit (Macherey-Nagel, Allentown, PA). Following viral RNA extraction, samples were evaluated using RT-qPCR to determine the viral RNA load in each animal37. Control standards (obtained from NIH AIDS Reagent Program) with known quantities of HIV-1 genome copies were used as amplification controls, as well as to stablish a standard curve that was used to determine the viral RNA load, based on the cycle threshold (Ct) value

